# RNA binding regulation is a new dimension in the type I IFN response

**DOI:** 10.1101/2025.05.30.656608

**Authors:** Louisa Iselin, Yana Demyanenko, Natasha Palmalux, Azman Embarc Buh, Wael Kamel, Peter Simmonds, Shabaz Mohammed, Alfredo Castello

**Author notes:** Correspondence &.

## Abstract

The type I interferon (IFN-I) response shapes the intracellular environment to supress virus infection. Historically, this remodelling has been linked to the transcriptional induction of interferon stimulated genes (ISGs). However, IFN-I-driven post-translational regulation of proteins already present in the cell remains relatively unexplored. Here, we profiled the activity of cellular RNA-binding proteins (RBPs), which are key players in antiviral immunity. Using RNA interactome capture (RIC), we identified hundreds of RBPs whose association with RNA is regulated by IFN-I (IR-RBPs). Among these IR-RBPs are both known antiviral proteins and novel candidates identified in this study through a knockdown screen. By modifying RIC to study IR-RBPs’ phosphorylation states, we identified several putative instances of IFN-I-driven phospho-regulation of RNA binding. We experimentally confirmed this phospho-driven regulation for MATR3. Altogether, our results reveal a new dimension of the cell’s antiviral programme, in which the cellular RNA-bound proteome is remodelled by IFN-I.

## INTRODUCTION

The type I IFN (IFN-I) response is central to antiviral innate immunity in vertebrates. IFN-I is produced by virus infected cells following sensing of pathogen associated molecular patterns (PAMPs) by pattern recognition receptors (PRRs). IFN-I binds to the IFNα receptor (IFNAR) on neighbouring cells, acting as an ‘early warning system’ by initiating a signalling cascade that prepares cells to combat infection. The canonical IFN-I-induced signalling cascade involves formation and nuclear translocation of the ISGF3 complex, which drives the upregulation of IFN-stimulated genes (ISGs) (Ivashkiv and Donlin 2014). Early microarray studies and more recent efforts using next generation sequencing have identified thousands of ISGs, across a wide range of cellular contexts (Der et al. 1998; de Veer et al. 2001; Rusinova et al. 2013; Shaw et al. 2017). ISGs can encode proteins that contribute to sensing and signalling or that directly restrict viral infection (Schoggins 2019). Several high-throughput screens have assessed the antiviral activity of putative ISGs, revealing broad-spectrum and virus-specific antiviral factors (Schoggins et al. 2011, 2014). However, genome-wide knockdown screens assessing antiviral activity following IFN-I treatment have found that many proteins modulating infection are not ISGs, speaking to additional layers of IFN-driven regulation that have yet to be explored (Zhao et al. 2012; Richardson et al. 2018).

There is mounting evidence that ISG induction is only one facet of the regulatory program induced by IFN-I. RNAseq studies have indicated that alternative splicing, alternative start site usage, and alternative polyadenylation are additional mechanisms that shape the transcriptome and, consequently, the proteome of IFN-I stimulated cells (Schneider et al. 2014; Colli et al. 2020; Ueda et al. 2024). Additionally, recent proteomic studies have reported discordances between RNA and protein abundance in the IFN-I response (Colli et al. 2020; Hodge et al. 2020; Chen et al. 2025). Indeed, a deep proteomic analysis of IFN-I treated HEK293 cells indicated that, of over 400 regulated mRNAs, only fewer than one hundred exhibited an equivalent change in protein abundance (Chen et al. 2025). This RNA/protein discordance hints at diverse levels of regulation occurring beyond transcript abundance.

In addition to transcriptional and post-transcriptional regulation, proteins can be post-translationally regulated in diverse ways. There is already extensive evidence for the essential role of protein post-translational modifications (PTMs) in canonical IFN-I signalling (Ivashkiv and Donlin 2014). For example, phosphorylation has been linked to signalling through RLR, TLR, and cGAS-STING pathways (Liu et al. 2015) and is central to signalling downstream of IFN-I/IFNAR interaction (Stark and Darnell 2012). In addition to this canonical role for phosphorylation, phospho-proteomic studies have indicated extensive modification of proteins not directly involved in IFN-I signalling (Viengkhou et al. 2021; Busnadiego et al. 2025; Chen et al. 2025). Beyond phosphorylation, many other modifications can play roles beyond signalling in the IFN-I response. For example, ISG15 is added to newly translated proteins with diverse functions following IFN-I response induction (Durfee et al. 2010). How these non-signalling modifications shape the intracellular environment has yet to be elucidated. However, there is an ever-expanding array of protein functions that are modulated by IFN-I, including localisation (Zuo et al. 2022), interactors (Kerr et al. 2020; Schiefer and Hale 2024), enzymatic activity (Kaur et al. 2008; Witteveldt et al. 2018), and stability (Zheng et al. 2011; Bhattacharya et al. 2014). Understanding how the IFN-I response is driving these changes will be an important area of study going forward.

One level of regulation that has not been comprehensively studied is the effect of IFN-I on cellular RBPs. RBPs are a heterogeneous group of proteins, classically identified through the presence of well-defined RNA-binding domains (RBDs) (Lunde et al. 2007). However, the complement of known RBPs, as well as our understanding of the diversity of RBPs’ function, has expanded dramatically over the past two decades. This has come with the development of new methods that comprehensively survey protein-RNA interactions, such as RNA interactome capture (RIC) (Baltz et al. 2012; Castello et al. 2012; Nechay and Kleiner 2020) . These approaches have uncovered hundreds of novel RBPs, many of which lack classical RBDs. Expansion of the RNA-bound proteome (RBPome) has led to a similar expansion in our understanding of the complexity of RNA-protein interaction and regulatory dynamics. For example, in addition to RBPs regulating RNA, it is now clear that RNA can act as a regulator of RBP activity – a process known as riboregulation (Hentze et al. 2018; Castello et al. 2024).

RBPs are central to antiviral innate immunity. Many PRRs are RBPs that bind to unusual features in viral (v)RNA (Schlee and Hartmann 2016). Additionally, many antiviral RBPs directly engage with vRNA to supress viral infection (Girardi et al. 2021). For example, IFIT1 binds to unmethylated cap structures found in some viral genomes and blocks translation of viral proteins (Kimura et al. 2013; Habjan et al. 2013; Fleith et al. 2018). Non-ISG RBPs have also been linked to innate immunity. These include the U2 small nuclear ribonucleoprotein (snRNP), which relocalises to the cytoplasm upon Sindbis virus (SINV) infection and blocks replication by engaging with the vRNA via a U2 snRNA-dependent mechanism (Kamel et al. 2024). Additionally, RBPs play broad roles in transcriptional and post-transcriptional regulation of ISG mRNAs. For example, ELAVL1 binds to ISG mRNAs following IRF3 stimulation, promoting their stability (Rothamel et al. 2021). RBPs are highly dynamic and can be regulated by numerous cellular stimuli, including viral infection (Garcia-Moreno et al. 2019; Kamel et al. 2021) and LPS stimulation (Liepelt et al. 2016). We hypothesised that IFN-I is another stimulus that can remodel the cellular RBPome, creating an intracellular environment that facilitates ISG expression and virus suppression.

In this study, we employed RIC to profile the dynamics and regulation of the cellular RBPome in response to IFN-I treatment. We thus defined the IFN-regulated (IR-)RBPome and, using an siRNA screen, identified novel players in antiviral immunity that are regulated exclusively at the level of RNA binding. To explore one factor that might shape the IR-RBPome, we extended RIC to include a phospho-enrichment step, allowing us to link changes in the IR-RBPs’ phosphorylation state with changes in RNA binding.

## RESULTS

### The RNA-bound proteome is regulated by IFN-I

We hypothesised that IFN-I signalling could globally remodel the RBPome to modulate the cellular susceptibility to infection. To test this possibility, we applied RNA interactome capture (RIC) to cells treated with IFNα2 for 0, 4, 8, 16 and 20 hours (Figure 1a). RIC employs ultraviolet (UV) light and denaturing oligo(dT) purification to specifically capture proteins bound to polyadenylated RNA (Castello et al. 2013). To assess the RNA-dependency of isolated proteins, in addition to UV irradiated samples, we included a mock, non-irradiated control. Silver staining of samples before LC-MS/MS yielded results consistent with prior RIC studies, with RIC eluate samples displaying a distinct banding pattern to input samples (Castello et al. 2012; Garcia-Moreno et al. 2019) (Figure S1a). Additionally, no proteins were detected in non-irradiated samples (beyond the RNases used in the processing of all samples), highlighting the strong UV-dependency of capture.

**Figure 1.**
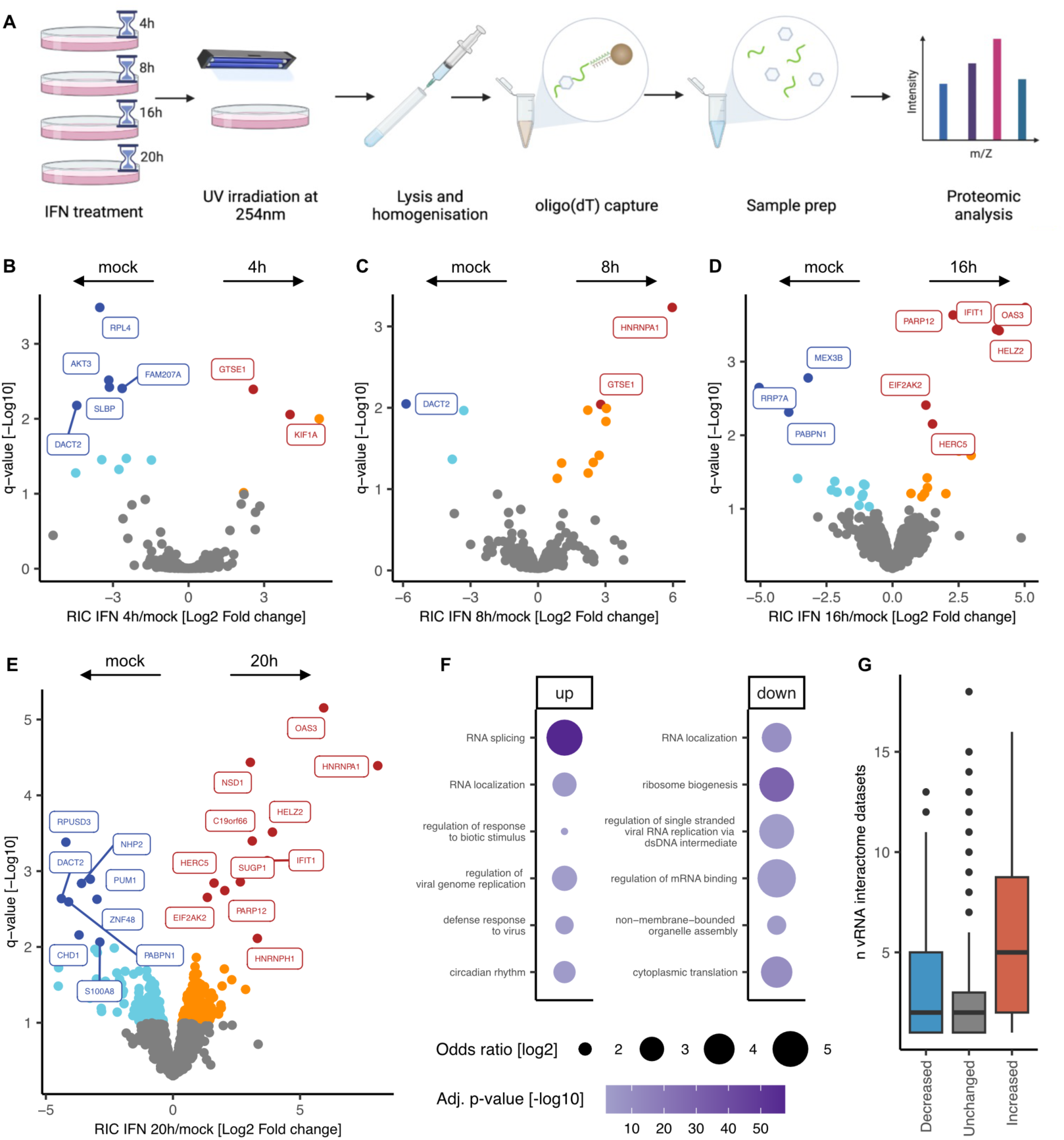
IFN-I remodels the cellular RNA-bound proteome. **(a)** Schematic representation of the IFN RIC protocol. **(b-e)** Volcano plots showing the log2 fold change and the false discovery rate (-log10 q-value) for each protein at 4, 8, 16 and 20 hours post IFN-I treatment, relative to mock. Data shown represents four biological replicates. Proteins with an FDR of 1% are labelled and are coloured dark red and dark blue. Proteins with an FDR of 10% are coloured orange and cyan. **(f)** GO enrichment analysis of biological process terms in proteins with significantly increased or decreased RNA-binding activity at 20h (10% FDR). **(g)** A boxplot indicating the number of vRNA interactome datasets that IR-RBPs and non-IR-RBPs occur in. vRNA interactome data from (Iselin et al. 2022).

By LC-MS/MS analysis, we identified and quantified a total of 1026 proteins across four replicates. All UV-irradiated samples exhibited high levels of consistency in raw intensity distributions (Figure S1b) and high correlation coefficients (Figure S1c), speaking to dataset quality. Quantitative analysis comparing each IFN-I treatment timepoint to untreated samples (0hpt), revealed subtle changes in the RBPome at 4 and 8hpt and more substantial remodelling at 16 and 20hpt (Figure 1b-e, Table S1). IFN-I-induced changes peaked at 20 hpt, with 150 stimulated and 122 inhibited RBPs (10% FDR). The time-dependent effects of IFN-I-driven RBPome remodelling were also observable by principal component analysis (PCA). In the absence of non-UV-irradiated samples, which otherwise account with the majority of variance in the dataset (Figure S1d), two distinct clusters of samples could be distinguished. These clusters corresponded to 0, 4 and 8h, and 16 and 20h post IFN-I treatment (Figure S1e). In summary, RIC uncovered over 250 proteins with altered RNA-binding activity following IFN-I treatment. We will henceforth refer to these as IFN-regulated (IR-)RBPs.

Gene ontology (GO) analysis identified distinct RNA binding related functions among stimulated and inhibited IR-RBPs (Figure 1f and S2a). Indeed, while more than 95% of all IR-RBPs had been previously identified as RNA interactors (Figure S2b), splicing related GO terms were highly but exclusively enriched in the stimulated RBP group. A more detailed analysis of these splicing factors revealed 8 SR proteins, 16 HNRNPs, cofactors of the U2 snRNP, and others. By contrast, splicing related proteins were absent in IFN-inhibited RBPs. This might reflect the central role of splicing regulation in shaping the antiviral cellular state (Liao and Garcia-Blanco 2021), or non-canonical roles for splicing regulators in innate immunity (Kamel et al. 2024).

Importantly, stimulated RBPs are also enriched in GO terms related to antiviral innate immunity (Figure 1F). Text mining to identify virus and immune-linked literature for each protein further confirmed the enrichment of antiviral activities among stimulated but not inhibited IR-RBPs (Figure S2c). Some of the proteins with the strongest increase in RNA binding at 20h were well-known antiviral proteins, such as OAS3, HELZ2, IFIT1, and PKR (EIF2AK2) (Schneider et al. 2014) at 1% FDR, and TRIM25 (Gack et al. 2007), TRIM56 (Wang et al. 2011), and IFI16 (Unterholzner et al. 2010) at 10% FDR. 48% of IFN-stimulated RBPs have links to immunity and 20% were previously related to IFN response. The excellent agreement between the current knowledge and our results strongly supports the quality of this dataset. Simultaneously, the substantial proportion of IR-RBPs that have not been linked to immunity offers new avenues for discovery.

Methods to study the vRNA interactome have now been applied to at least 11 viruses from 6 viral families, offering a rich resource for identifying cellular RBPs that bind to vRNA (Iselin et al. 2022; Castello et al. 2024). By compiling these datasets, we previously defined a core vRNA interactome of proteins that interact with the RNA of coronaviruses, flaviviruses and alphaviruses. IFN-stimulated RBPs are enriched in core vRNA interactome members, reflecting strong links with viral infection (Figure S1d) (Iselin et al. 2022). No enrichment is observed among IFN-inhibited RBPs, although a subset of these downregulated IR-RBPs occur in a large number of vRNA interactomes (Figure 1g). Manual inspection highlighted that many of these IFN-inhibited RBPs have established roles in promoting virus infection, including XRN1, PATL1, UPF1, PURA, PURB, and MKRN2. It is plausible that IFN-I not only stimulates RBPs with antiviral roles but may also reduce the activity of proteins that are required to sustain infection.

We next examined the regulatory patterns of all IR-RBPs over the course of the IFN-I response using k-means clustering. We identified six RNA-binding activity clusters: i) early and strong induction; ii) early but mild induction; iii) late induction; iv) early inhibition followed by recovery; v) early inhibition; and vi) late inhibition (Figure 2a, Table S2). ISGs appeared in two clusters (i) early/strong up and ii) early/mild up), suggesting differences in activation kinetics (Figure 2b). Interestingly, other functionally related groups of proteins also had matching kinetic profiles. For example, all annotated P-body associated proteins exhibit decreased RNA-binding activity, with PATL1, UPF1, RC3H2, and APOBEC3F following similar patterns of inhibition (Figure 2c). Additionally, core paraspeckle proteins followed a strikingly consistent activation-inhibition-activation pattern (Figure 2d). To test whether non-core, paraspeckle associated proteins follow the same trend, we used a recent meta-analysis of paraspeckle composition that includes 84 proteins (McCluggage and Fox 2021). Of these 84 proteins, 34 had increased RNA-binding activity upon IFN treatment, while only 4 exhibited decreased RNA binding (Figures 2e). Strikingly, over 60% of these stimulated IR-RBPs displayed patterns of induction similar to core paraspeckle proteins (Figures 2e&f). These findings suggest that the IFN-I response modulates paraspeckle dynamics in a highly controlled manner, calling for further experiments to elucidate how this regulation relates to known or novel roles of paraspeckles and their component parts in innate immunity (Girardi et al. 2023; Milcamps and and Michiels 2024). These time-resolved IR-RBP kinetics are a valuable tool for understanding the regulatory dynamics of individual proteins and protein networks.

**Figure 2.**
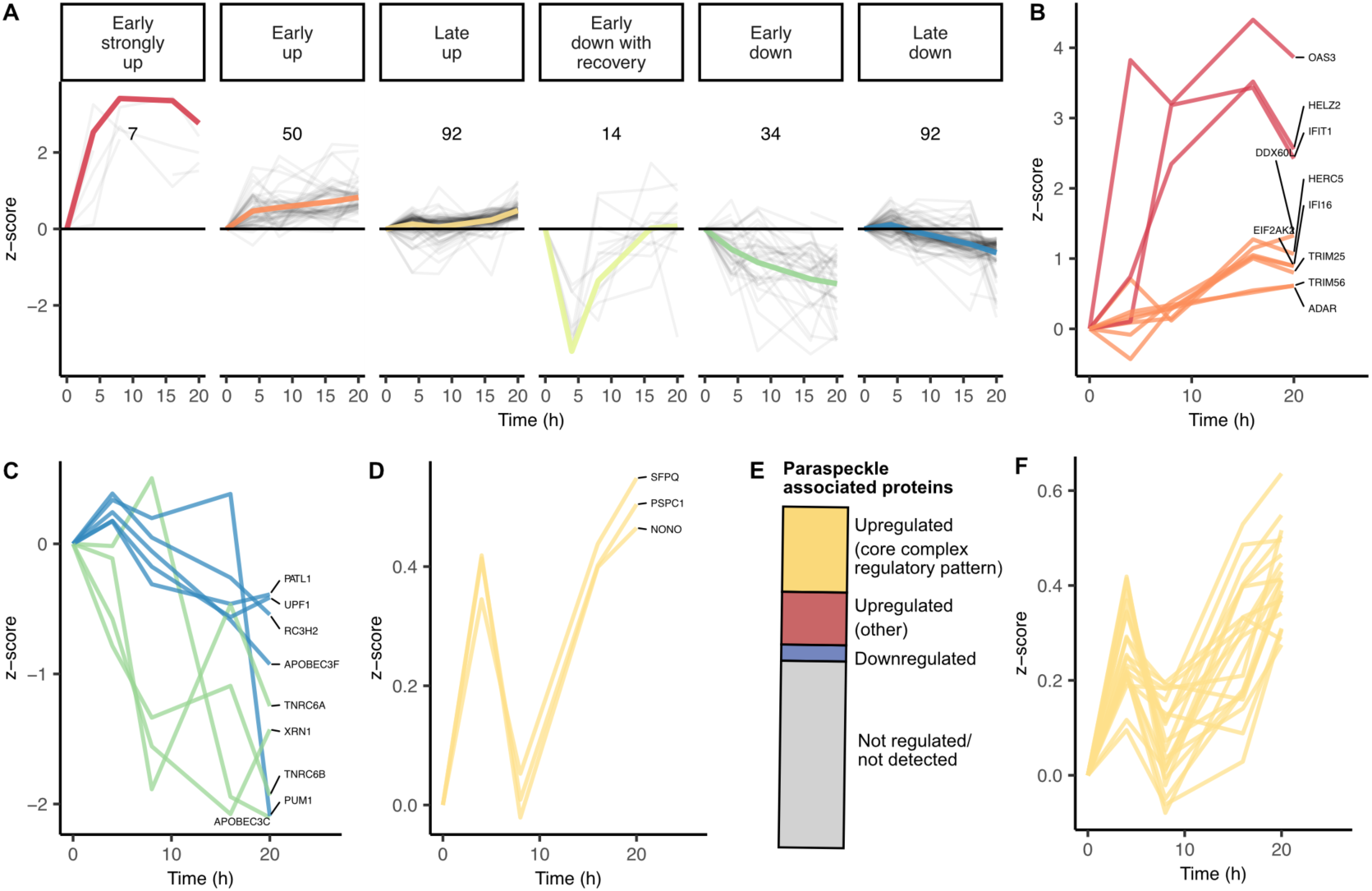
IR-RBPs exhibit distinct kinetic profiles in response to IFN-I. **(a)** K-means clustering results of significantly changing RBPs’ z-scores across all four timepoints. Thick coloured lines represent the mean z-score profile for each cluster. Profiles for individual proteins are shown in grey. Numbers on each plot represent cluster size. **(b-d)** Kinetic profiles of all proteins classified as ISGs (b), P-body associated (c), or core paraspeckle components (d). **(e)** Classification of paraspeckle associated proteins (defined by (McCluggage and Fox 2021)), based on their behaviour in the IFN-I RIC dataset. **(f)** Kinetic profiles of all significantly regulated, paraspeckle associated RBPs that display the core complex regulatory pattern.

### The IR-RBPome is regulated independently of protein abundance

Differential association of RBPs with RNA can be driven by changes in protein abundance, substrate RNA abundance, protein localisation, or through post-translational modification (Mukhopadhyay et al. 2009). IFN-I stimulation can alter the expression of mRNAs and proteins (e.g. ISGs)). To determine whether the observed RNA-binding changes are mediated by matching alterations in protein abundance, we compared our results to deep transcriptomic and proteomic data generated in the same cell line and under the same experimental conditions in our laboratory (Chen et al. 2025). Strikingly, only 9 proteins exhibit significant changes in RNA, protein and RNA-binding levels, while the majority of IR-RBPs were regulated exclusively at the level of RNA binding (Figure 3a). Comparison of the fold changes in the RIC dataset versus the whole cell proteome and transcriptome confirmed the notable discordance between RNA-binding activity and protein and mRNA levels (R=0.25 and 0.19, respectively) (Figure 3b-c). Together, these findings suggest that the regulation of most IR-RBPs upon IFN-I treatment occurs at the level of RNA-binding activity rather than through changes in transcript or protein abundance.

**Figure 3.**
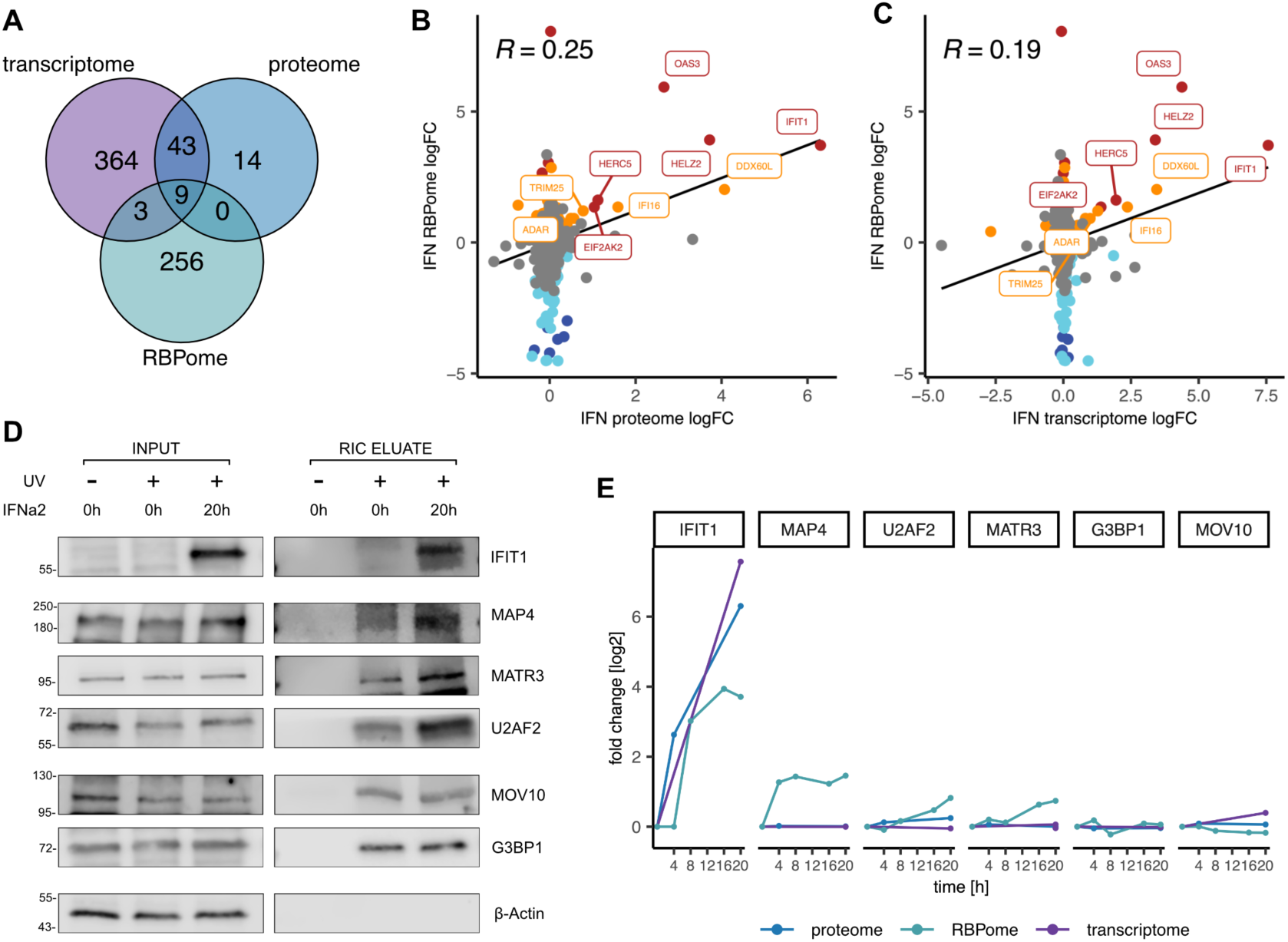
Changes in RNA binding are not due to changes in protein abundance. **(a)** Venn diagram comparing significant hits in the transcriptome (5% FDR), proteome (10% FDR), and RBPome (10% FDR) 20 hours post IFN-I treatment. Transcriptome and proteome data taken from (Chen et al. 2025). **(b&c)** Scatter plots comparing the fold change (log2) in the RBPome versus the proteome (b) and transcriptome (c). Points are coloured based on significance in RIC dataset. Pearson correlation coefficients are displayed in the top left of the plots. **(d)** Western blots showing relative changes in RIC inputs and eluates. **(e)** Kinetic profiles of changes in transcriptome, proteome and RBPome of each protein assessed by western blot in panel d.

To validate this striking discordance, we generated RIC input (whole cell lysate) and eluate (RNA bound fraction) samples for analysis by Western blot. This orthogonal analysis revealed an IFN-I-dependent increase in IFIT1 levels in both input and eluates, which is consistent with an abundance-driven effect, as observed in the proteomics data (Figure 3d-e). On the other hand, the IR-RBPs MATR3, MAP4, and U2AF2 increased their interaction with RNA without equivalent changes in protein abundance (Figure 3d-e). The association with RNA of non-IR-RBPs G3BP1 and MOV10 was not affected by IFN-I and, as expected, the negative control β-Actin was not detected in eluates. Altogether, these results suggest a model in which IFN-I regulates the activity of RBPs independently of protein abundance.

### IR-RBPs are novel players in antiviral immunity

RBPs with IFN-enhanced RNA-binding activity may represent a new class of antiviral proteins, analogous to ISGs. Complementarily, IFN-mediated inhibition of RBPs required for infection could be an alternative strategy to restrict infection. To test these hypotheses, we performed a targeted siRNA screen with 13 proteins for their effect on Sindbis virus (SINV) infection. SINV is a model alphavirus that is known to induce an IFN-I response in HEK293 cells (Garcia-Moreno et al. 2019) and for which the vRNA interactome has been well-established (Kamel et al. 2024). Candidates were selected based on: i) pattern of induction, ii) available literature reflecting possible roles in viral infection, and iii) their occurrence in vRNA interactomes (Iselin et al. 2022). We reasoned that proteins that interact with the RNA of different viruses are more likely to be involved in infection. Details of each selected candidate are outlined in Table S3.

To assess the roles of these proteins in infection, we incubated HEK293 cells with specific siRNAs for 48h, then infected them with SINV for a further 18h (Figure S3a). The efficiency of siRNA knockdown was assessed using RT-qPCR, with most proteins (bar RBM39 and PNUTS) showing an mRNA depletion of at least a 50% (Figure 4a). Despite achieving only partial depletion, 10 out of 13 of the candidates tested resulted in a significant change in SINV vRNA levels, with six of these leading to at least a doubling or halving viral gene expression (Figure 4b). Knockdown of 9 candidates also led to significantly altered capsid protein levels, measured by Western blotting (Figures 4c&d). Of these, knockdown of ATXN2, JUP, RALY, THRAP3, SUGP1, U2AF2, PNUTS, MAP4 and AKAP8 led to increased viral infection, suggesting that these proteins may contribute to the antiviral response. On the other hand, RAVER1 and GIGYF2 knockdown led to decreased infection, suggesting that these are required for SINV infection. The high frequency of hits in this screen speaks to the importance of IR-RBPs in shaping viral infection outcomes, and to the richness of the IFN RIC dataset as a resource for identifying novel players in infection.

**Figure 4.**
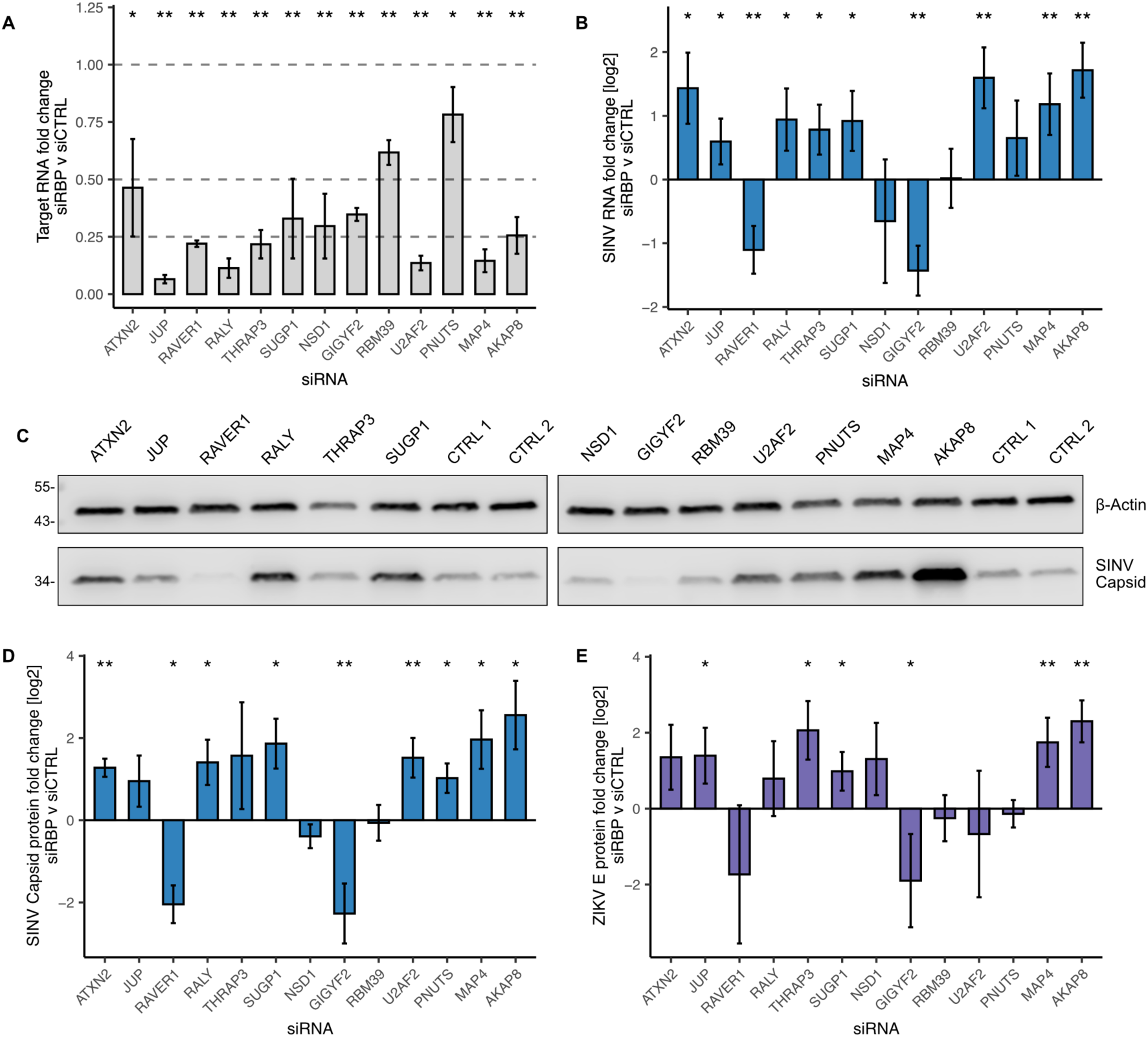
A targeted screen of IR-RBPs highlights novel players in RNA virus infection. **(a)** RT-qPCR analysis of knockdown efficiency of siRNAs targeting 13 IR-RBP candidates 48h post transfection. Target RNA levels were normalised to 18S ribosomal RNA and fold change was calculated relative to the average expression in two siRNA controls. Individual values are a mean of technical duplicates, while bars represent the mean of at least four biological replicates ± SD. Significance was calculated using the one-sample t-test, following confirmation of data normality by the Shapiro-Wilks test. **(b)** RT-qPCR analysis of the effect of IR-RBP knockdown on SINV RNA levels. Fold change and significance were calculated as described for (a). Values are a mean of technical triplicates and at least four biological replicates were included per siRNA. **(c)** Representative Western blot of the effect of 48h siRNA knockdown of 13 IR-RBP candidates on SINV capsid protein levels. Control siRNA treated samples (CTRL 1 and CTRL 2) were loaded on both membranes. **(d)** Quantification of the protein signal for at least four biological replicates from western blot analysis shown in (c). Quantification represents the log2 fold change of β-actin-normalised capsid levels, relative to the average of two siRNA controls. Significance was calculated as in (a). **(e)** Western blot quantification results showing the effect of siRNA knockdown of the 13 IR-RBP candidates on ZIKV E protein levels. A representative blot is shown in Figure S3. Quantification represents the log2 fold change of β-actin-normalised E protein levels, relative to an siRNA control. Significance was calculated as in (a). P<0.01 ** ; P<0.05 *.

The IFN-I response has evolved to respond to diverse pathogens. To determine whether the effects observed in SINV were conserved against other RNA viruses, we extended the screen to zika virus (ZIKV, flavivirus genus) (Figure S3b). After 24h of ZIKV infection in knocked down cells, we observed six proteins whose knockdown significantly altered ZIKV infection (Figure 4e and S3c). These included SUGP1, GIGYF2, MAP4 and AKAP8, which also had significant effects on SINV infection, suggesting that these IR-RBPs play conserved roles in the antiviral immune response against different families of RNA viruses.

### IFN-regulated phosphorylation can drive changes in RNA-binding activity

Given the widespread activity-level regulation of RBPs in response to IFN-I, we hypothesised that protein post-translation modifications (PTMs) could be important in shaping the IR-RBPome. However, methods to globally assess how PTMs alter RBP activity are lacking. Here, we developed a novel approach, termed phospho-RIC, to determine how the altered phosphorylation state of RBPs in response to a stimulus relate to their RNA-binding activity. IFN RIC eluates were subjected to phosphopeptide enrichment using Zr-IMAC and analysed by LC-MS/MS. Phospho-RIC thus enables the measurement of the abundance of a given phosphosite in RNA-bound protein across different experimental conditions and allows us to directly compare the level of phosphorylation with the level of RNA-binding activity. We identified an average of 961 phosphosites in UV irradiated samples, while in non-irradiated controls, an average of 173 phosphosites were detected (Figure S4b). Approximately 68% of the peptide intensity signal in UV crosslinked samples corresponded to phosphopeptides, indicating a successful enrichment (Figure S4c).

Many phosphosites exhibited altered abundance at 4h, 8h and/or 20h (Figure 5a and S4d-e, Table S4). Since both RIC and phospho-RIC datasets were generated from the same samples, we can directly compare RBPs’ RNA-binding activity with their phosphorylation status (Figure 5b). Many of the significantly changing phosphosites occurred either in proteins that weren’t reliably detected in the original IFN RIC experiment or proteins that do not show significantly altered RNA-binding activity, suggesting these phosphosites are not modulating interaction with RNA (Figure 5c). Excitingly, there was also a subset of significantly changing phosphosites that occurred in IR-RBPs. Some of these changes matched the observed changes in overall RNA-binding activity (i.e. more protein bound, more phosphosite), preventing us from drawing inferences about their functional relevance. However, we also observed instances of antagonistic behaviour between phosphosite abundance and RNA binding (i.e. more binding and less phosphosite or vice versa) (Figure 5d-j).

**Figure 5.**
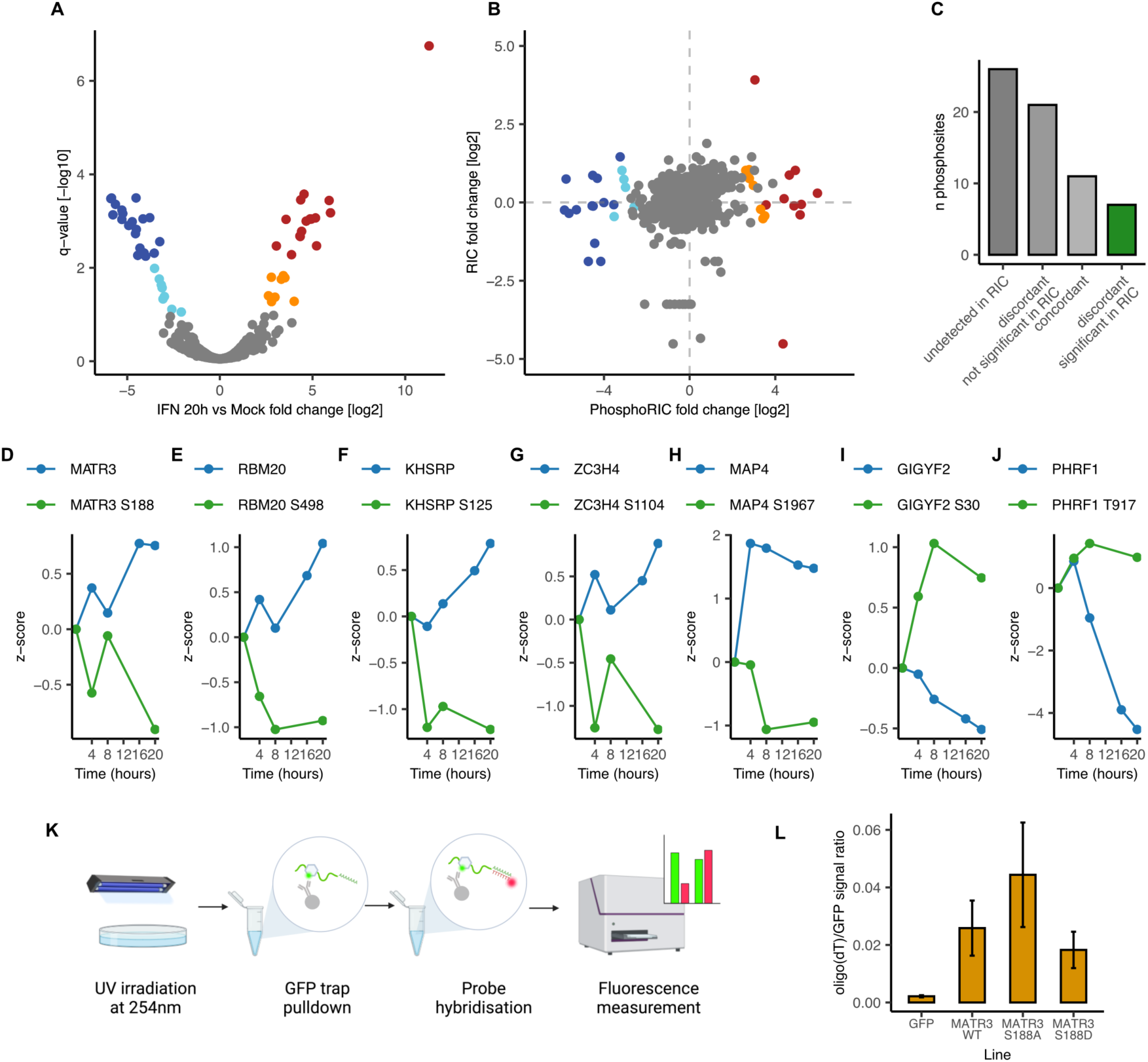
Defining the IFN-I-regulated RNA-bound phosphoproteome. **(a)** Volcano plot showing the log2 fold change and the false discovery rate (-log10 q-value) for each identified phosphopeptide at 20 hours post IFN-I treatment, relative to mock. Peptides with an FDR of 1% are labelled and are coloured dark red and dark blue. Peptides with an FDR of 10% are coloured orange and cyan. Data shown represents three biological replicates. **(b)** A scatter plot comparing the log2 fold change in phosphopeptide abundance with the log2 fold change in whole protein abundance in the RIC dataset (Figure 1e) at 20h post IFN-I treatment. Points are coloured based on significance in the phospho-RIC dataset. **(c)** Counts of significant hits in the phospho-RIC dataset, according to the binding profiles of proteins in the original RIC dataset. **(d-j)** Kinetic profiles of z-scores in phospho-RIC (green) and RIC (blue) datasets for RBPs with discordant phosphosites. **(k)** Schematic of the dual fluorescence RNA binding assay. **(l)** Ratio of oligo(dT) (Alexa Fluor 594) to GFP signal, reflecting RNA-binding level for GFP, MATR3-GFP, MATR3-GFP S188A, and MATR3-GFP S188D. All intensity values were blank corrected. Bars represent mean ratio across three biological replicates ± SD.

Because of the negative charge of the phosphate group, phosphorylation can act as a negative regulator of protein-RNA interactions, causing repulsion effects with the phosphate backbone of the RNA. Indeed, proteome-wide analysis of RNA-binding domains (RBDs) revealed a high level of correlation between phosphorylation and other post-translational modifications and RNA-binding surfaces (Castello et al. 2016). Therefore, we hypothesised that these antagonistic phosphosites could be contributing to proteins’ altered RNA-binding activity in response to IFN-I. We tested this hypothesis for one such regulated RBP: MATR3 (Figure 5d).

MATR3 has pleiotropic effects on RNA metabolism, with roles ranging from alternative splicing regulation to DNA damage response, translation control, and beyond (Coelho et al. 2015; Sprunger and Jackrel 2024). MATR3 has also been found to translocate to the cytoplasm in SINV and coxsackievirus B3 (CVB3) infected cells and play an antiviral role (Kamel et al. 2024). Experimental evidence pinpointed the high-confidence RBDs of MATR3 to its RRMs and an upstream intrinsically disordered region (IDR) (Castello et al. 2016). The S188 phosphosite maps to this RNA-binding IDR (Figure S4f). To test if S188 phosphorylation regulates MATR3 RNA-binding activity, we generated HEK293 FITR lines expressing either wild-type protein tagged with GFP, a phospho-mimetic mutant (S188D), or a phospho-null mutant (S188A). Using these lines, we assessed MATR3 RNA-binding using an *in cellulo* dual-fluorescence RNA-binding assay (Castello et al. 2012; Strein et al. 2014). In brief, HEK293 FITR cells expressing a GFP fusion RBP are crosslinked with UV, followed by the immunoprecipitation of the covalently linked protein-RNA complexes with GFP-trap. Following stringent washes, oligo(dT) probes labelled with a red fluorophore (Alexa 594) are hybridised to RBP-associated RNA. The ratio of red to green fluorescence represents RNA/protein signal (Figure 5k). This assay showed a clear trend towards the phospho-null mutant having higher RNA-binding activity relative to WT MATR3 or the phospho-mimetic mutant (Figure 5L & Figures S4G-H). These results are consistent with the IFN RIC/ phospho-RIC analysis that suggested that IFN-induced dephosphorylation of S188 increases MATR3 RNA-binding activity.

To further characterise this regulatory phosphosite, we performed iCLIP2 on WT and mutant MATR3.This approach employs *in cellulo* UV crosslinking, immunoprecipitation and library preparation to generate a genome-wide map of the binding sites of a protein with single nucleotide resolution (Buchbender et al. 2020). Interestingly, the S188A mutant had more significant binding sites and target mRNAs than the WT protein, while the S188D mutation had fewer (Figure 6a). These results confirmed orthogonally the findings with the dual fluorescence assay, further demonstrating that MATR3 RNA-binding activity is enhanced when the IDR is not phosphorylated. Strikingly, the target mRNAs and the binding sites within them were conserved in WT and phospho-mutants, with WT and S188D’s smaller target sets representing a subset of S188A’s targets (Figure 6b). The biotype of the target mRNAs was preserved across MATR3 variants, with a strong preference for binding to mRNAs (Figure 6c). MATR3 and its mutants bind predominantly to introns, consistent with MATR3’s known role in splicing regulation (Figure 6d) (Uemura et al. 2017). Motif enrichment analysis revealed the same pyrimidine-rich sequence motif in all cases (Figure 6e), which reflects the known binding preference of MATR3. Moreover, the incidence of these motifs within binding sites was similar across MATR3 variants (Figure 6f). These results indicate that MATR3’s sequence specificity is mainly driven by its RRMs, while the IDR in which S188 sits is involved in tuning RNA binding affinity. Phosphorylation of S188 thus causes a reduction in MATR3’s affinity for RNA, which is relieved upon IFNα treatment due to S188 dephosphorylation.

**Figure 6.**
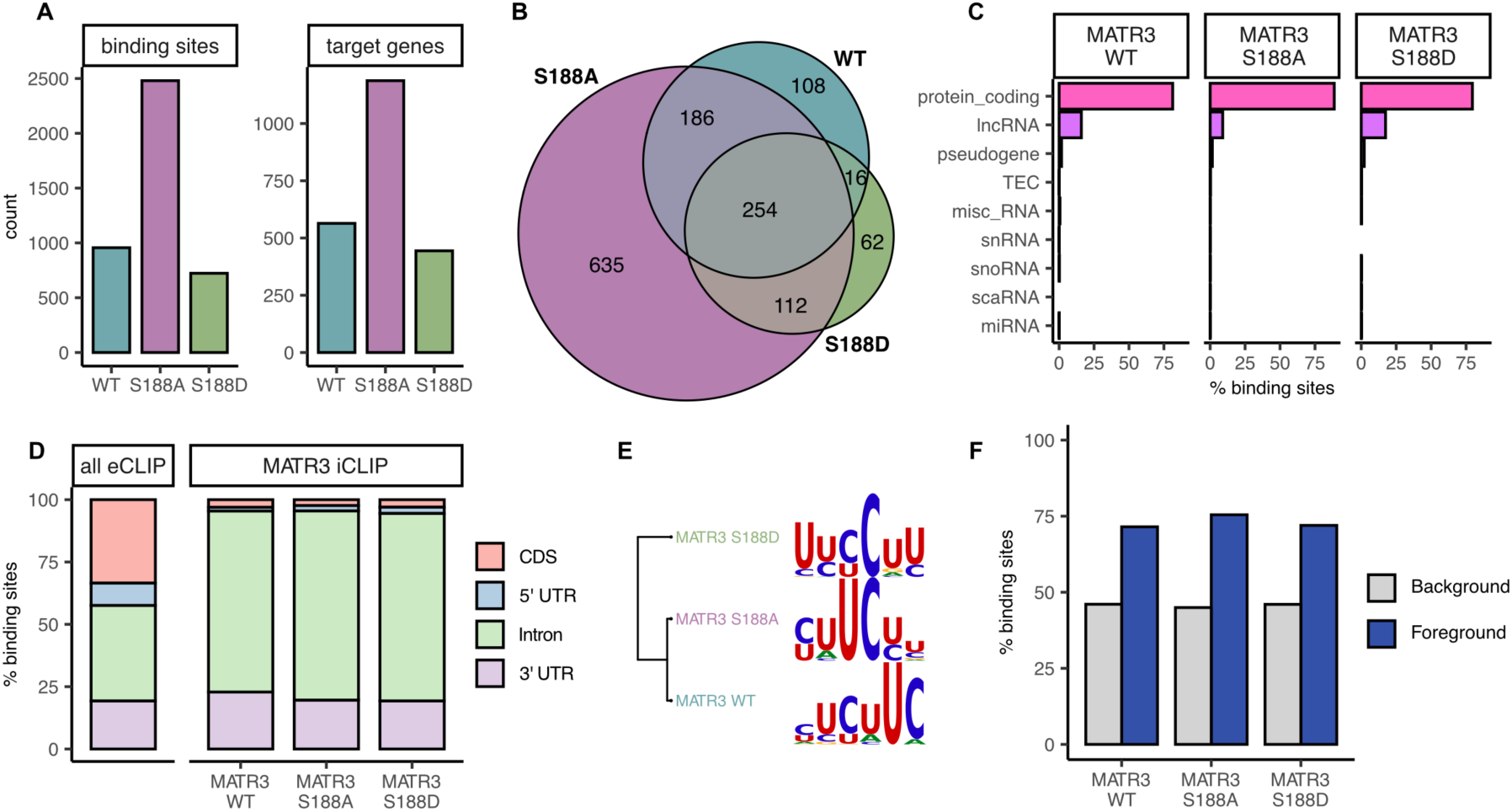
MATR3’s affinity, but not specificity, is modulated by S188 phosphorylation. **(a)** Bar plots showing the number of binding sites and target genes in mock samples from WT, S188A, and S188D MATR3-GFP. **(b)** Venn diagram showing overlap between WT and mutant MATR3 target genes. Target genes are defined as genes with at least one significant binding site, based on DEW-seq enrichment analysis over the SMI control (adj. p-value < 0.01). **(c)** Bar plots showing the proportion of target RNAs corresponding to different RNA biotypes bound by each MATR3 variant. **(d)** Stacked bar plot showing the distribution of significantly enriched binding sites in protein coding mRNAs within each region of the RNA, based on genome annotation. A composite of all eCLIP datasets from the ENCODE database is included as a reference. **(e)** Hierarchical clustering of the most enriched motif in each of the three datasets. Motif enrichment was performed on 50 nucleotide windows centred around the signal peak of each binding site. Enrichment was relative to gene and gene region-matched sequences generated for each binding site and was performed using STREME from the MEME suite (Bailey et al. 2015). **(f)** Percentage of sequences containing at least one motif in MATR3 binding sites (blue bar) compared to the gene and gene region matched control sequences.

## DISCUSSION

The type I IFN response is a key aspect of antiviral innate immunity. Hitherto, it has been primarily studied as a transcriptional response. However, there is mounting evidence that IFN-I shapes the cellular landscape in diverse ways beyond ISG induction. Here, we characterise one such regulatory facet: the modulation of RNA-binding activity of RBPs. Despite the known central roles of RBPs in viral sensing and restriction, there has yet to be a global effort to assess whether, and which, RBPs are regulated in response to IFN-I signalling. Using RIC, we identified over 250 RBPs exhibiting altered RNA-binding activity. This dataset sheds light on a new regulatory dimension in the IFN-I response and represents a valuable resource in defining a novel set of IFN-regulated proteins.

Several IR-RBPs are ISGs that are known to serve central roles in the antiviral immune response. The identification of the pattern recognition receptors OAS3, PKR, IFIT1 and IFI16 in our study was surprising, given their known roles in specifically detecting vRNA (Schlee and Hartmann 2016). However, there is a growing body of evidence for the importance of cellular RNA in PRR regulation (Gokhale et al. 2024) and for PRRs as regulators of cellular mRNA. For example, PKR binds to a stem loop in the 5’ UTR of the IFNƔ mRNA, repressing its translation (Cohen-Chalamish et al. 2009). An exciting area of future investigation will be to establish whether equivalent regulatory mechanisms could account for the other PRRs’ cellular binding activity and, if so, what their RNA targets might be.

In addition to stimulated IR-RBPs that might have analogous antiviral functions to ISGs, around half of IR-RBPs exhibit decreased RNA-binding activity. An exciting question is what the purpose of downregulating over one hundred RBPs in response to IFN-I might be. Strikingly, we found amongst down-regulated RBPs many instances of host proteins known to be co-opted by diverse viruses to promote infection. For example, MKRN2 was recently shown to be important for influenza A virus (IAV) and SINV (Bonazza et al. 2024; Kamel et al. 2024), and PURA and B play central roles in HIV RNA metabolism (Garcia-Moreno et al. 2023). The exonuclease XRN1 and its cofactor PATL1 are other IFN-downregulated RBPs that are essential for virus infection (Pijlman et al. 2008; Scheller et al. 2009; Garcia-Moreno et al. 2019; Liu et al. 2021; Ruscica et al. 2024). For example, XRN1 mediates widespread degradation of cellular mRNAs upon alphavirus infection that increases the availability of nucleotides required for vRNA replication (Ruscica et al. 2024). Additionally, in flavivirus infection, XRN1-driven vRNA degradation gives rise to a highly structured sfRNA that has been linked to innate immune evasion (Pijlman et al. 2008). Given their diverse and critical roles in infection, it is plausible that XRN1 and PATL1’s downregulation by IFN-I could be a means of restricting viral replication.

As well as reducing availability of host dependency factors, downregulation of IR-RBPs could also be involved in de-repressing IFN-I signalling. For example, GIGYF2 is a negative regulator of IFNβ mRNA translation, whose inhibitory activity is enhanced through the activity of SARS-CoV2 Nsp2 (Xu et al. 2022). Therefore, GIGYF2’s reduced RNA-binding activity following IFN-I treatment might be a means of promoting IFN-I signal propagation by reducing its interaction with the IFNβ mRNA. Indeed, we found that GIGYF2 knockdown represses both SINV and ZIKV, which is consistent with a higher capacity to synthesise IFNβ in the absence of this RBP. Further exploration of these downregulated IR-RBPs will be required to determine whether others play similar roles in either promoting viral infection or inhibiting IFN-I signalling.

Two ISGs, APOBEC3C (A3C) and APOBEC3F (A3F), were unexpectedly among the downregulated IR-RBPs (Peng et al. 2006; Bonvin et al. 2006). APOBEC3 proteins induce C to U editing in cellular and viral nucleic acids, which can be antiviral by inducing sequence changes in the viral genome. While A3C and A3F exclusively edit ssDNA (Pecori et al. 2022), RNA binding is central to their regulation. They are packaged into large RNP complexes as soon as they are translated and are sequestered within P-bodies and stress granules under resting conditions (Smith 2016). Their interaction with RNA is thought to directly inhibit their editing ability, preventing damaging mutations of cellular DNA (Gallois-Montbrun et al. 2007). Upon infection, APOBEC3s are released from these complexes, allowing them to engage with viral nucleic acids. A3C and A3F might similarly be released from repressive RNP complexes following IFN-I treatment, explaining their decreased RNA-binding activity. This reflects the complexity of RNA-protein interactions and the variety of regulatory dynamics that can exist at this interface. Moreover, it highlights the value of an unbiased and comprehensive survey of the RBPome in discovering new regulatory processes even for established players in antiviral immunity.

In addition to identifying known players in viral infection, we also uncovered novel antiviral candidates through a small-scale, targeted screen. Out of the 13 proteins screened, nine exhibited significant antiviral activity in SINV infection despite siRNAs only mediating partial knockdowns. These results suggest that beyond focusing on ISGs, there is value in considering proteins that are regulated at the “activity level” in response to IFN-I. However, it is important to note that the changes that we observe are relatively mild (2-4 fold repression) compared to dominant antiviral factors such as PKR. The differences in repression penetrance between these and canonical antiviral factor could be related to technical factors, such as incomplete knockdown or functional redundancy that would require combinatorial knockdowns to address. Alternatively, it could reflect differences in biological roles, with ISGs being the “mediators” of the antiviral responses and IR-RBPs contributing to “modulation” or fine-tuning activity. For example, IR-RBPs could exert roles as ISG’s accessory factors or might be involved in regulating the response itself, as in the case of GIGYF2.

Here, we performed a targeted screen as a proof of concept that antiviral RBPs could be identified within this pool of proteins. However, we only tested the effects of 13 proteins. We hope that future research will extend our efforts to cover a wider range of IR-RBPs. Additionally, we only focused on two RNA viruses. These represent a fraction of the diverse challenges that the IFN-I response has evolved to respond to, which include other classes of viruses, other pathogens (McNab et al. 2015), and cancer (Holicek et al. 2024). Understanding which combinations of IR-RBPs are involved in shaping the response to these diverse threats represents another important avenue of future research.

As well as highlighting the value of this dataset, the siRNA screen also identified exciting candidates in their own rights, each of which warrants further investigation. For example, MAP4 is a microtubule associated protein and non-canonical RBP (Castello et al. 2016) that we previously identified as an intriguing member of the core vRNA interactome (Iselin et al. 2022). In this study, we found that its RNA-binding activity is strongly enhanced by IFN-I and that it exhibits a pronounced antiviral effect in both SINV and ZIKV infection. Additionally, we observed that its phosphorylation at S822 anticorrelates with its association with RNA. S822 maps to an IDR that overlaps with both the RNA-binding and microtubule binding (Chapin and Bulinski 1991) regions of MAP4. Understanding the contribution of MAP4’s RNA binding, microtubule binding and phosphorylation state to its antiviral activity will shed light on the complex regulatory processes that moonlighting antiviral proteins undergo in response to IFN-I stimulation. Another interesting candidate that had conserved antiviral effects on SINV and ZIKV is AKAP8 (also known as AKAP95). AKAP8 is a protein kinase A (PKA) adaptor, which acts as a scaffold in PKA nuclear microdomains (Clister et al. 2019). One possibility is that AKAP8 is involved in coordinating the changes in phosphorylation landscape that help in shaping the antiviral state.

An important finding of this study was that most IR-RBPs do not exhibit altered protein abundance, raising questions as to what is responsible for their altered RNA binding. Changes in RNA substrate abundance can be an important determinant of RBP regulation. This is known to be a driver during SINV infection, where the transcriptome is pervasively remodelled (Garcia-Moreno et al. 2019). While IFN-I also causes changes to the transcriptome, these are relatively subtle and very specific when compared to the effects observed in SINV infection (Chen et al. 2025). Therefore, while it is plausible that certain IR-RBPs are regulated by changes in mRNA substrate abundance, it is unlikely that “substrate availability” is the cause of the regulation of over 250 RBPs. RNA binding can also be modulated by changes in protein localisation, interaction partners, or post-translational modifications (Sampath et al. 2003). We hypothesise that all these factors contribute to a greater or lesser extent in the regulation IR-RBPs.

In this study, we assessed the contribution of phosphorylation, which is a critical post-translational modification in IFN-I signalling (Zhou et al. 2017). RBPs can be heavily modified by phosphorylation (England et al. 2022) and, due to its physicochemical properties, the phosphate group can act as a powerful regulator of RNA-binding activity (Velázquez-Cruz et al. 2021). However, establishing the regulatory significance of specific phosphosites is a substantial challenge in the field of phospho-proteomics (Needham et al. 2019). A recent study developed an elegant approach (qRIC) to assess the effect of phosphosites on overall RIC pulldown efficiency under steady state conditions through quantitative comparison with whole cell lysate (Vieira-Vieira et al. 2022). This approach highlighted the value of linking phosphorylation information with RNA-binding activity data. In our approach, we focus on comparing phosphosite abundance and RNA-binding activity dynamics in response to a stimulus. In this way, we can link phosphosite behaviour with a direct readout for RNA-binding activity and with a context in which phospho-regulation might be involved. This allowed us to rapidly generate testable hypotheses, which we experimentally validated for MATR3. This is a powerful extension of RIC that could be applied to diverse cellular contexts to explore phospho-driven regulatory dynamics following perturbation. Additionally, by combining RIC with other PTM enrichment protocols, or even with iterative enrichment for multiple PTMs, it would be possible to more comprehensively assess the PTM regulatory landscape in the IFN-I response and beyond. Given the scarcity of examples of RBPs whose mechanism of regulation in response to a given stimulus is understood, this has the potential to enhance our understanding of post-translational regulation of RNA-binding activity in general and of specific RBPs of interest.

## LIMITATIONS OF THIS STUDY

RIC is a powerful tool for identifying proteins associated with poly(A) RNA (Baltz et al. 2012; Castello et al. 2012; Hentze et al. 2018). However, it is not without its biases. As with any proteomic based study, RIC is affected by protein abundance and length, as well as trypsin and MS peptide compatibility. Additionally, UV_254_ crosslinking is known to be biased towards specific nucleotides with U being the most efficient. UV_254_ crosslinking is also benefited by the duration and prevalence of the interaction, and by the existence of zero distance contacts between the protein’s amino acids and the nucleotide bases (Iselin et al. 2022). While UV_254_ crosslinking is very specific, it is less efficient for protein-RNA interactions that exclusively involve the ribose-phosphate backbone of the RNA. This is a potential limitation to detect proteins that bind preferentially to double-stranded RNA, many of which are important in the antiviral response. However, PKR (EIF2K2) and ADAR can still be identified in RIC studies (Iselin et al. 2022), suggesting that this limitation does not apply equally to all dsRNA-binding proteins. Another bias in RIC is the focus on polyadenylated RNA (mainly mRNA) because of the oligo(dT) capture step (Castello et al. 2012). Therefore, proteins binding to non-poly(A) non-coding RNAs, which are known to play roles in innate immunity (Schneider et al. 2014), will not be captured by RIC. Therefore, complementing RIC with complementary methods that captures bulk RNA would be a valuable approach. However, it is important to keep in mind that rRNA and tRNAs are the most abundant nucleic acids in total RNA isolation approaches. Because of their different biases, both approaches are complementary rather redundant.

The phospho-RIC approach requires an additional purification step with the phospho-enrichment that causes sample loss. Therefore, the amount of input material is critical for deep phospho-RBPome surveys. In this proof-of-principle experiment we used ∼20x10^6^, but we anticipate that the depth of the analysis would be greater using 5-10x more cells. Therefore, phospho-RIC application to precious and limited samples such as biopsies or low growth primary cells, might be challenging.

## DATA AVAILABILITY

The mass spectrometry proteomics data have been deposited to the ProteomeXchange Consortium via the PRIDE (Perez-Riverol et al. 2025) partner repository with the dataset identifier PXD064360. iCLIP sequencing raw data has been submitted to the GEO repository with the dataset identifier GSE294181.

## ACKNOWLEDGEMENTS

A. C. is funded by the European Research Council (ERC) Consolidator Grant ‘‘vRNP-capture’’ 101001634 and the MRC grants MR/R021562/1 and MC_UU_00034/2. L.I. was funded by the BBSRC DTP scholarship number BB/M011224/1. S. M and Y.D. are funded by an EPSRC grant (V011359/1 (P)). N.P. was funded by the MRC CVR PhD programme. W.K. was funded by the European Union’s Horizon 2020 Marie-Sklodowska-Curie fellowship 842067. A.E.B. was funded by Fundación Ramon Areces postdoctoral fellowship program.

We thank past and present members of the Castello and Mohammed groups for scientific discussion in the development of this manuscript. We thank Honglin Chen, Jeff Lee and Aino Jarvelin for support in developing analysis scripts. We thank Arvind Patel for ZIKV E antibody.

While this work is funded by the European Union, views and opinions expressed are however those of the author(s) only and do not necessarily reflect those of the European Union or the European Research Council Executive Agency. Neither the European Union nor the granting authority can be held responsible for them.

## AUTHOR CONTRIBUTIONS

Conceptualization, L.I., W.K., S.M., A.C.; Methodology, L.I., Y.D., W.K., S.M., A.C.; Investigation, L.I., Y.D., A.E-B., N.P.; Writing, original draft, L.I., A.C.; Writing, editing, L.I., A.E-B., N.P., Y.D., W.K., A.C, S.M., P.S.; Funding acquisition, A.C., S.M.; Resources, A.C., S.M.; Supervision, A.C, S.M., P.S., W.K.

## MATERIALS AND METHODS

### Cell culture

Cells were grown at 37°C with 5% CO_2_ and maintained in Dulbecco’s modified eagle medium (DMEM; Gibco, 41965039), with 10% fetal bovine serum (FBS; Gibco, 10500064) and 1x penicillin-streptomycin (Sigma Aldrich, P4458). The media of HEK293 Flp/In T-REx (FITR) parental line (Thermo Fisher Scientific # R78007) was supplemented with 100 µg/ml Zeocin (Gibco, #R25001) and 5 µg/ml Blasticidin S Hydrochloride (Thermo Fisher Scientific, J67216.8EQ). HEK293 FITR overexpression lines were kept in 150 µg/ml Hygromycin B (Invitrogen, #10687010) and 5 µg/ml Blasticidin S Hydrochloride.

To generate HEK293 FITR overexpression lines, cells were transfected with pOG44 and pcDNA5-FTR-TO (containing gene of interest) using X-tremeGENE 360 transfection reagent (Roche, XTG360-RO), as per manufacturer’s instructions. Cells were selected with 150 µg/ml Hygromycin B. Overexpression was induced by supplementing media with 1 µg/ml doxycycline and was confirmed by Western blotting with specific antibodies. Mutant constructs were generated by PCR mutagenesis, using the pcDNA5-RBP-FTR-TO plasmid as template and primers designed with Agilent’s QuikChange Primer Design tool (sequences in Table S6). PCR was performed with PFU turbo DNA polymerase (Agilent, #600252) and the template plasmid was degraded using DpnI (NEB, R0176S). Mutations were confirmed by Sanger sequencing.

### Viruses

Sindbis virus (pT7-SVwt (Castelló et al. 2006)) and Zika virus (ZIKV-WT MR766 - ATCC VR-84 strain) were produced by *in vitro* transcription from the plasmid, followed by transfection of transcribed vRNA into BHK21 cells using Lipofectamine 3000 (Invitrogen, 11668019). Virus titration was performed by plaque assay in VeroE6 cells.

### siRNA knockdowns

1x10^5^ HEK293 FITR parental cells per well or 1.25x10^4^ A546 cells were seeded in a 24 well plate 24 hours prior to transfection. 5 pmol of each siRNA (Thermo Fisher, Silencer Select, siRNA sequences in table S6) was transfected using Lipofectamine RNAiMAX (Invitrogen, L3000001). HEK293 FITR cells were infected with WT SINV (MOI 0.1) after 48 hours and harvested 18 hours post infection (hpi). A549 cells were infected with WT ZIKV (MOI 0.2) for 24h prior to harvesting. For harvesting, cells were washed once in cold PBS and lysed either in RIPA buffer (10 mM Tris HCl pH 7.5, 150 mM NaCl, 0.5 mM EDTA, 0.1% SDS, 1% Triton X-100, 1% sodium deoxycholate) for western blotting analysis or in RLT buffer for RT-qPCR analysis (Qiagen, 79216).

### Conventional protein analysis

Samples were boiled in NuPAGE 4x LDS sample buffer pfrior to gel loading (Invitrogen, NP0007). Proteins were then separated by SDS-PAGE on 4-20% polyacrylamide gels (BioRad, 4561095). Silver staining was performed using the SilverQuest silver staining kit, according to their basic staining protocol (Invitrogen, LC6070). Western blotting was performed as previously described in (Garcia-Moreno et al. 2019). Membranes were imaged on the LI-COR Odyssey Fc or LI-COR Odyssey CLx. Details of antibodies used can be found in Table S6.

### RT-qPCR

Cells were harvested in RLT buffer and RNA was extracted using the RNeasy Mini Kit (Qiagen, 74104). Reverse transcription and RT-qPCR analysis was performed by Luna universal one-step RT-qPCR kit (NEB, #E3005L) with the primers listed in Table S6.

### RNA interactome capture

#### IFN RIC

Two 15cm dishes per condition were seeded with 20x10^6^ HEK293 cells per dish in DMEM (10% FBS, 1x P/S). After 24 hours, 4hr, 8hr, 16hr, and 20hr plates were treated with 200U/ml recombinant human IFNα 2a (Novus Biologicals, NBP2-34971). At 4, 8, 16, and 20 hours after IFN treatment, the plates for each timepoint were crosslinked and harvested. One plate from mock crosslinked and non-crosslinked controls was harvested at 4 hours and the other was harvested at 20 hours (end of the experiment). To crosslink, plates were washed twice in cold PBS and UV irradiated at 254nm at an energy of 150mJ/cm^2^ on ice. Cells were then harvested using a cell scraper in 5ml RIC lysis buffer (20 mM Tris-HCl pH 7.5, 500 mM LiCl, 0.5% LiDS wt/vol, 1 mM EDTA, 0.1% IGEPAL (NP-40) and 5 mM DTT) per condition and were stored at -80°C. Lysates were homogenised by repeated passage through a 5ml syringe with 27G needle until clear. 300µL/ 15cm dish of pre-calibrated oligo(dT)_25_ magnetic beads (NEB, S1419S) were added to each sample and the RIC protocol was performed as outlined in (Garcia-Moreno et al. 2019). Two consecutive rounds of capture were performed. Samples were eluted in 360 μL elution buffer and were then RNase treated with RNase T1 and RNase A (1:1 in 10-fold dilution with elution buffer, 1μL per sample) at 37°C for 1 hour.

For proteomic analysis, samples were processed using the bead-based single-pot, solid-phase-enhanced sample-preparation (SP3) method (Hughes et al. 2019). First, samples were incubated with 10mM TCEP and 50mM C-IAA at RT in the dark for 30 minutes to reduce and alkylate proteins. Next, samples were split to ensure maximum volume per tube of 300 μL and 100μg SpeedBeads Magnetic Carboxylate Modified Particles (Sigma-Aldrich, GE651521 05050250) were added to each tube. 100% ACN was added to achieve a final ACN concentration of 80% and samples were incubated for 30 minutes at RT. Samples were then washed twice with 70% EtOH and once with 100% ACN in a loop until they were free from detergent. Samples were incubated overnight at 37°C with 250ng Trypsin Gold (Mass Spec Grade) (Promega, V5280). The next day, supernatant was transferred to fresh tubes and residual peptides left on the beads were eluted in 2% DMSO for 2 minutes off the rack. Eluates were combined and acidified with neat formic acid.

For western blot analysis, samples were processed as above up to the RNase digestion step, with two 15cm dishes (20x10^6^ HEK293 cells/dish) used per condition. RIC eluates were concentrated using Amicon Ultra-0.5 3kDa Centrifugal filter (Millipore, UFC500308) prior to loading for western blot.

#### Phosphoenrichment

Following trypsin digestion, the peptide mixture was treated with benzonase nuclease (1:1000) and incubated at 37◦C for 30 min. Samples were then cleaned up using the Oasis HLB 96-well μElution Plate (Waters, 186001829), as per manufacturer’s instructions. Peptides were eluted twice in 25 μL SPE elution buffer. For phosphoenrichment, 0.4mg of equilibrated MagReSyn Zr-IMAC beads were added to each sample (Resyn Biosciences, MR-ZRM002). Samples were incubated for 20 minutes (RT, 1350 RPM), then washed once in 400 μL Loading Buffer, twice in 400 μL Wash Buffer 1, and twice in 400 μL Wash Buffer 2. For each wash, samples were incubated for 2 min (RT, 1350 RPM), before being placed on the magnetic rack. Samples were incubated for 10 min (RT, 1350 RPM) in 1% ammonium hydroxide to elute peptides. Elution was repeated twice more, and eluates were acidified with formic acid to a final concentration of 4%. Clean-up was then repeated, as detailed above. Samples were eluted once in 25 μL SPE elution buffer and then diluted in 100 μL 5% DMSO 5% FA before loading onto the Mass Spectrometer.

#### Mass Spectrometry

Analysis of IFN RIC peptides was carried out using an Ultimate 3000 nano-LC 1000 system coupled to an Orbitrap Exploris (Thermo Fisher Scientific). Phosphoenriched peptides were analysed using an Ultimate 3000 nano-LC 1000 system coupled to an Orbitrap Fusion Lumos Tribrid Mass Spectrometer (Thermo Fisher Scientific). Peptides were initially trapped on a C18 PepMap100 pre-column (300 μm inner diameter x 5 mm, 100A) and then separated on an in-house built C18 column (Reprosil-Gold, Dr. Maisch, 1.9 μm particle size) column (ID: 75 μm, length: 50 cm) at a flow rate of 200 nL/min. IFN RIC peptides were separated over 60 min (12-35%B) and phospho-enriched peptides were separated over 30 min (12-38% B). In both cases, mobile phase A (water and 0.1% formic acid) and mobile phase B (acetonitrile and 0.1% formic acid) were used. Separated peptides were directly electrosprayed into the mass spectrometer. Mass spectra were acquired in the orbitrap (350-1400 m/z, resolution 60000, AGC target 3 x 106, maximum injection time 50 ms) in a data-dependent mode. The top 40 most abundant peaks in the survey scan were fragmented using HCD (resolution 7500, AGC target 4 x104, maximum injection time 64 ms).

### Dual fluorescence RNA binding assay

HEK293 FITR MATR3-eGFP (WT, S188A, and S188D) and HEK293 FITR eGFP lines were seeded in 6 well plates at a density of 1x10^6^ cells per well and induced with 1ug/ml doxycyline for 24 hours. Cells were then crosslinked and lysed in DF lysis buffer (500mM KCl, 5mM MgCl2, 10mM Tris pH 7.5, 1% IGEPAL, 1mM DTT), and the dual fluorescence assay was performed using the ChromoTek GFP-Trap Multiwell Plate (Chromotek, gtp). The protocol was performed as described in (Strein et al., 2014) with the following modifications. Probe hybridisation was performed in 50 μL DF hybridisation buffer (500mM LiCl, 1mM EDTA, 20mM Tris HCl pH 7.5, 0.01% IGEPAL, 0.05% LiDS, 5mM DTT), supplemented with 160nM oligo(dT)_25_ stellaris probe (Alexa Fluor 594) (Life technologies Ltd). Fluorescence measurements were taken using the BMG Clariostar platereader. Measurements were taken for GFP (470-15 absorption/ 515-20 emission) and Alexa Fluor 594 (575-20 absorption/630-40 emission). Each well was analysed as a 10x10 matrix in ‘well scan’ mode and the median fluorescence was calculated. Enhanced dynamic range was used and focal height was set using auto-focus, with measurements taken from above. All values were normalised to a ’blank’ well, in which no IP had been performed.

### iCLIP2

HEK293 FITR MATR3-eGFP (WT, S188A, and S188D) lines were seeded in 10cm dishes (10x10^6^) cells per dish in DMEM (5% FBS) supplemented with 1ug/ml Doxycycline. Cells were harvested after 24 hours and iCLIP2 was performed, as described in (Garcia-Moreno et al. 2023). Samples were quantified using Qubit DNA HS and library size was measured using High Sensitivity D1000 TapeStation. IP and SMI sample groups was pooled equimolarly and then mixed at the following proportions: 75% IP library pool and 25% SMI library pool. Sequencing was performed on a NextSeq 550 sequencer with a 75 cycle High-output kit v2.5 (Illumina, #20024906).

### Data analysis

#### IFN RIC and IFN phospho-RIC

Protein identification and quantification were performed using Andromeda search engine implemented in MaxQuant (Cox and Mann 2008). Peptides were searched v Human Uniprot database. FDR was set at 1% for both peptide and protein identification and ‘match between runs’ was turned on. Otherwise, default parameters were used.

For IFN RIC analysis, data analysis of ProteinGroups file from MaxQuant was performed in R. Data was normalised using the ‘vsn’ package. Rows were filtered to remove any with >2 NA values in each condition under study. Minimum value imputation was performed for on/off changes (all NA values in one condition and <2 NA values in the other). Only values for replicates corresponding to non-NA values in the other condition were imputed. Fold-changes and p values were calculated using the limma package (Smyth 2005; Ritchie et al. 2015), with replicate included as a block in the regression analysis, and FDR (q-value) was calculated from p-values using the fdrtool package. For IFN phospho-RIC samples, the same analysis pipeline was applied to phosphopeptides from the Phospho(STY) file.

Principal component analyses were performed using the base R package on log2 transformed values. Data was filtered to exclude N/A values and normalised and batch corrected to match limma analysis processing where appropriate.

For kinetic clustering and plotting, fold-change values were taken from limma analysis results where possible. For conditions where there were insufficient values for a protein to be included in the analysis, minimum imputation was performed and fold-changes were calculated relative to mock. Values were then transformed from fold-changes into z-scores. K-means clustering was performed using the FKM function from the fclust package with k (number of clusters) set to 6.

#### IFN RIC dataset characterisation

GO enrichment analyses were performed using clusterProfiler in R (Wu et al. 2021). A p-value cut-off of 0.01 and a q-value cut-off of 0.05 were used, and the Benjamini-Hochberg method was used to correct for multiple testing. Non-unique GO terms were collapsed using clusterProfiler’s simplify function.

Text mining was performed in R using the RISmed package. Separate searches of PubMed titles and abstracts were done for the terms: virus-linked ([gene name] AND [virus OR viral]), immune-linked ([gene name] AND [immune OR immunity]), and IFN-linked ([gene name] AND [IFN OR interferon]. No minimum date was set, and the maximum date was set to the date of analysis. A gene was considered ‘linked’ to the above terms if there were 6 or more papers returned.

#### iCLIP

ICLIP data processing and downstream analysis were performed as described in (Álvarez et al. 2024)

---

**Supplementary Figure 1.**
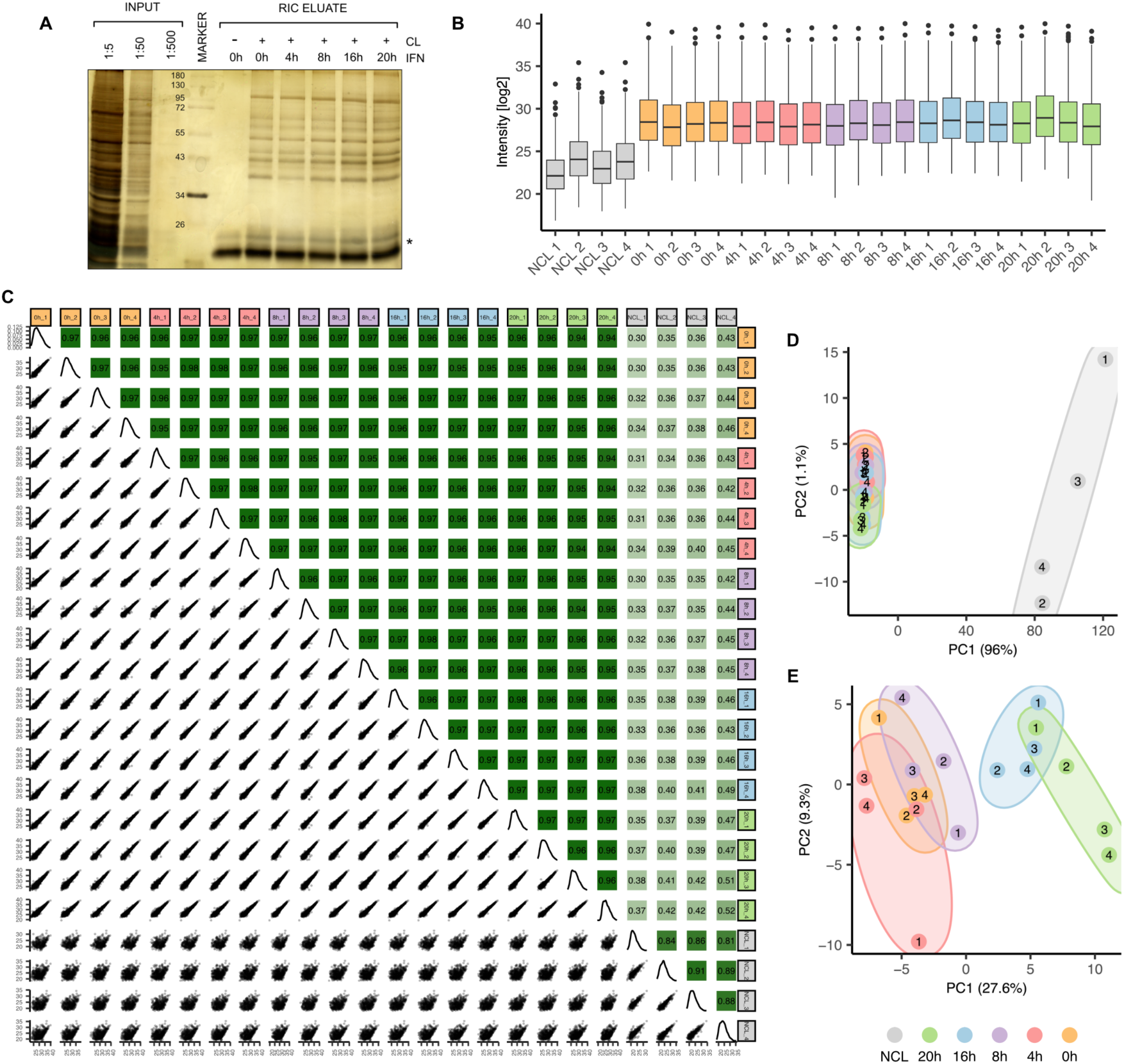
**a)** Silver staining analysis of input and RIC eluate samples. The * marks the bands produced by RNase A and RNase T1. Representative of all four biological replicates. **b)** Boxplot of LC-MS/MS log2 intensity distributions for IFN RIC samples prior to normalisation. **c)** Correlation analysis across all IFN RIC samples. Bottom: Scatter plots of log2 intensity values. Top: Pearson correlation coefficients, with boxes coloured to reflect strength of correlation. **d&e)** Principal component analysis (PCA) with (d) or without (e) non-crosslinked control samples included. Percent of variation explained by each principal component is indicated in brackets within the axis titles.

**Supplementary Figure 2.**
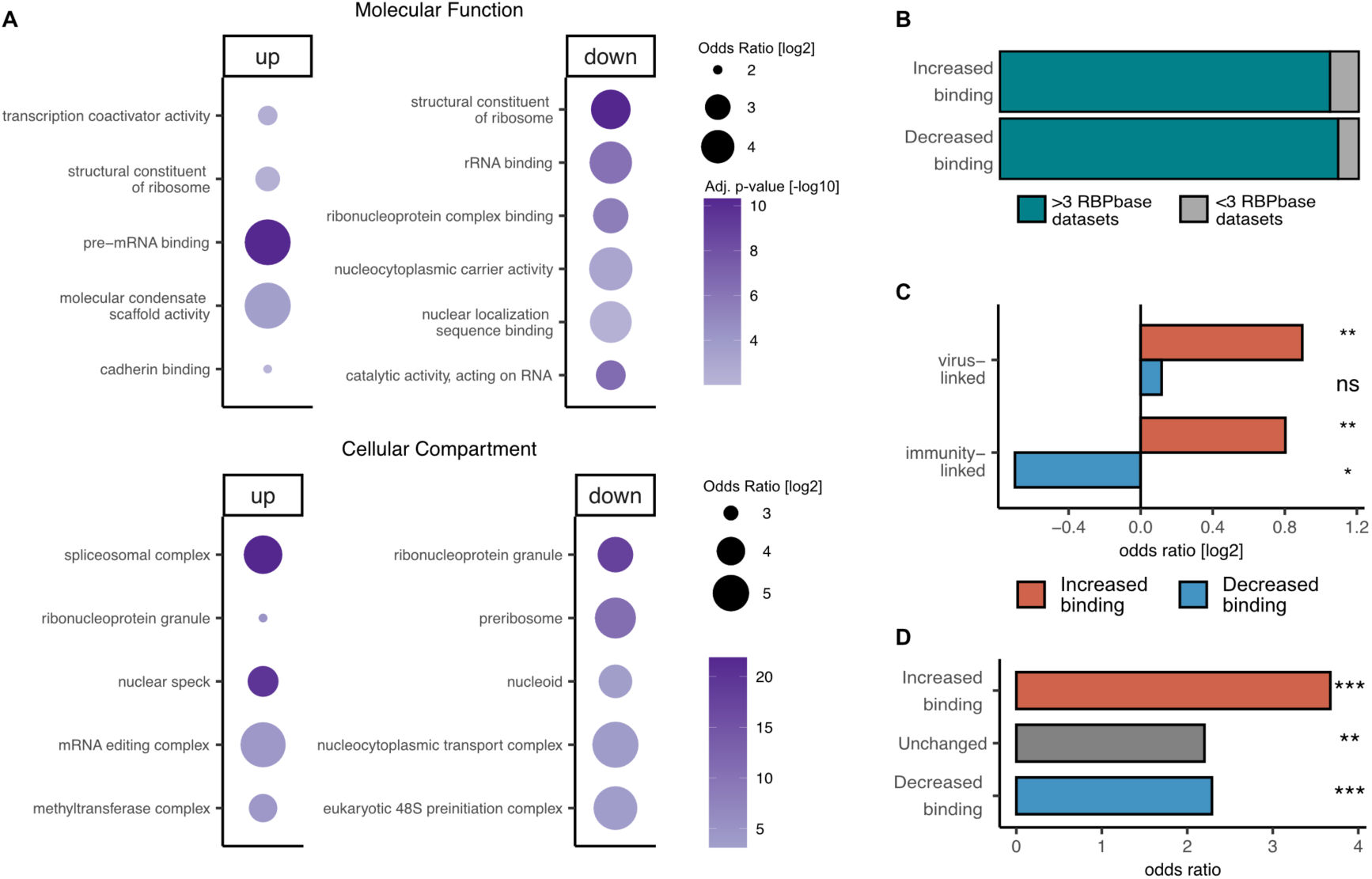
**a)** GO enrichment analysis of molecular function (top) and cellular component (bottom) terms in proteins with significantly increased or decreased RNA-binding activity at 20h (10% FDR). **b)** Proportion of significantly increased or decreased RBPs at 20h (10% FDR) that are known RBPs. Proteins were classified as RBPs if they occurred in three or more datasets in RBPbase. **c)** Enrichment analysis of significantly increased or decreased RBPs at 20h (10% FDR) linked to either immunity or viruses in the literature. A protein is considered ‘linked’ if there are 6 or more papers with the protein’s name and either ‘virus’/‘viral’ or ‘immunity’/‘immune’ mentioned in the title or abstract in PubMed. Statistical enrichment was assessed using Fisher’s Exact Test. **d)** Bar plot showing enrichment of proteins in the IFN RIC datasets in members of the core vRNA interactome (Iselin et al, 2022). Enrichment was tested using Fisher’s Exact Tests for proteins with increased, decreased, or unchanged RNA-binding activity at 20h. Significance: P<0.0001 *** ; P<0.01 ** ; P<0.05 *.

**Supplementary Figure 3.**
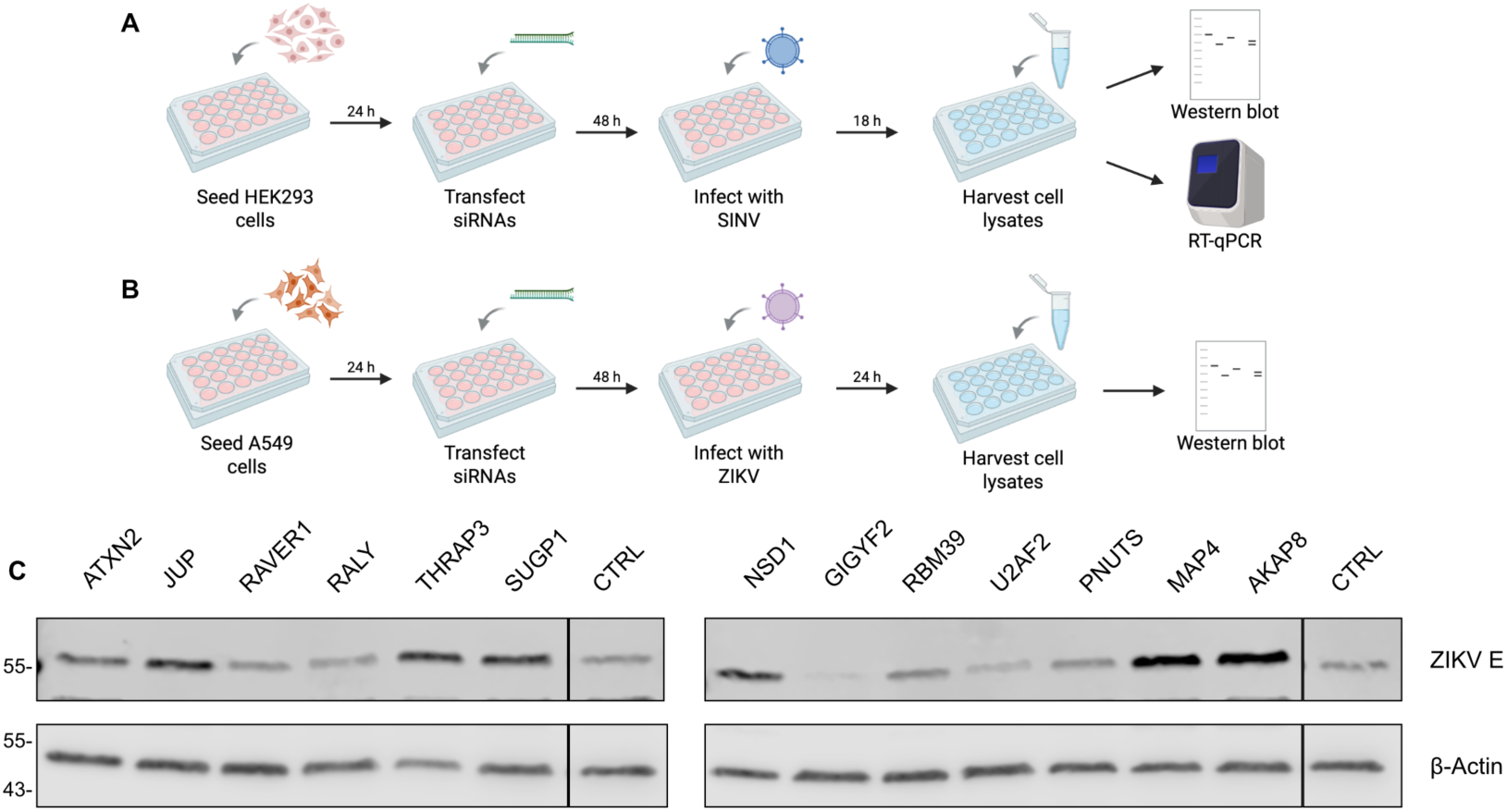
**a&b)** Schematic of the protocol used to screen IR-RBPs’ effect on SINV (a) and ZIKV (b) infection using siRNA knockdown. **c)** Representative western blot of the effect of 48h siRNA knockdown of 13 IR-RBP candidates on ZIKV E protein levels, divided across two membranes. Control siRNA treated sample was loaded on both membranes.

**Supplementary Figure 4.**
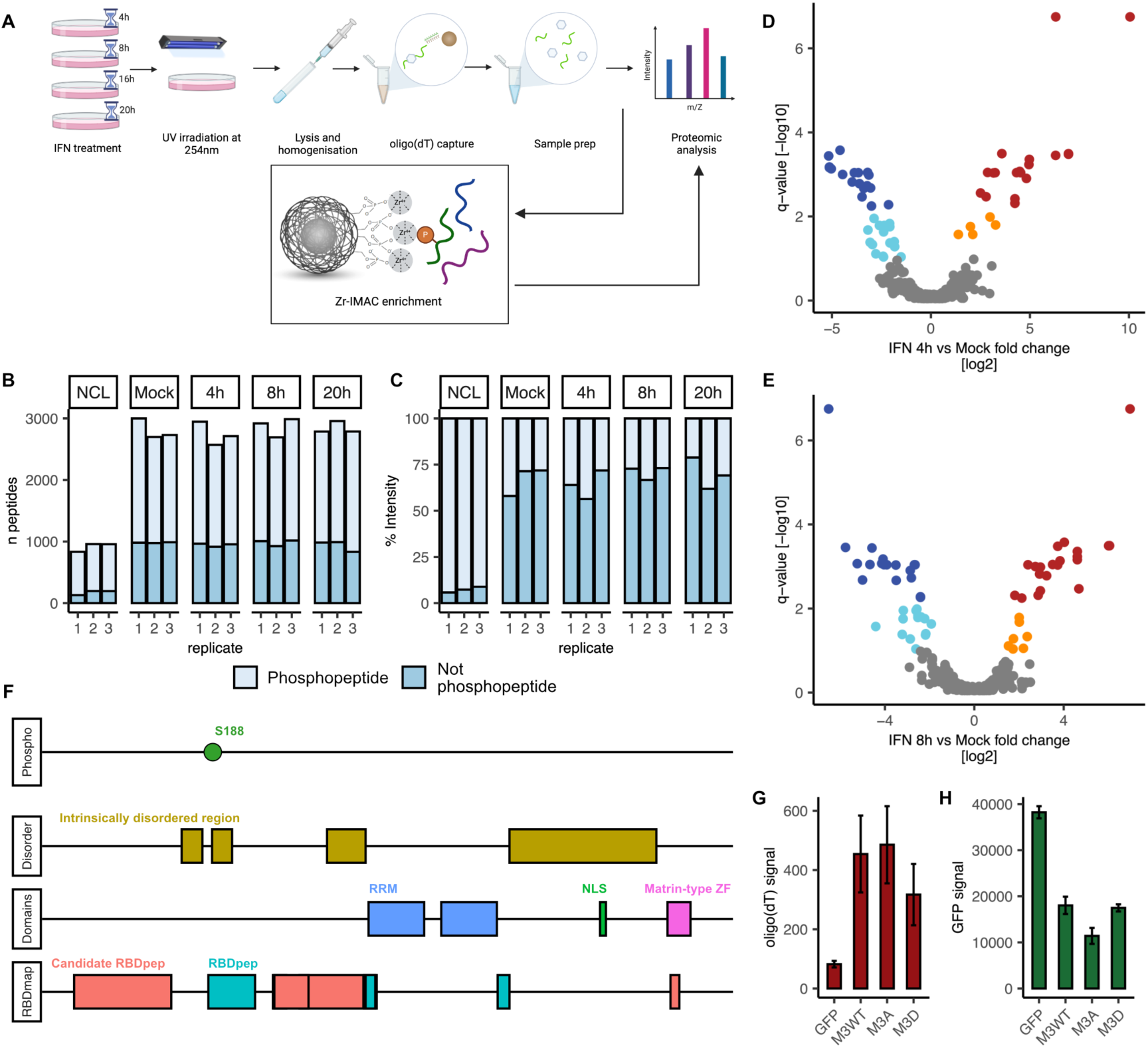
**a)** Schematic of phospho-RIC. **b)** Bar plot showing the number of peptides identified by LC-MS/MS for each sample. Number of phosphorylated and non-phosphorylated peptides are indicated by darker and lighter shades of blue, respectively. **c)** Bar plot indicating the proportion of overall signal coming from phosphorylated or non-phosphorylated peptides in each sample. **d&e)** Volcano plots showing the log2 fold change and the false discovery rate (-log10 q-value) for each identified phosphopeptide 4 hours (d) and 8 hours (e) post IFN-I treatment, relative to mock. N = 3. f) A schematic of MATR3’s pertinent features. (Phospho) Position of the IFN-regulated phosphosite. (Disorder) Regions annotated in Uniprot as disordered. (Domain) Pfam annotated domains (RBDmap) RNA-bound peptides identified at 1% FDR (RBDpep) and 10% FDR (Candidate RBDpep) are highlighted in turquoise and pink. g&h) Blank-corrected oligo(dT) (Alexa Fluor 594) (g) and GFP (h) signal from the RNA-binding dual-fluorescence assay. Bars represent mean values ± SD. N = 3.

